# AAV-mediated overexpression of Prdm12 in knee-innervating afferents reduces inflammatory joint pain and neuronal hyperexcitability in mice

**DOI:** 10.1101/2025.10.14.680517

**Authors:** Maya Dannawi, Luke A. Pattison, Alexander Cloake, Eric J. Bellefroid, Ewan St. John Smith

**Affiliations:** Department of Pharmacology, University of Cambridge, United Kingdom; ULB Neuroscience Institute, Université Libre de Bruxelles, Belgium

**Keywords:** Prdm12, Pain, Behavior, Electrophysiology, Transcription regulation, Gene Therapy

## Abstract

Inflammatory joint pain features in numerous musculoskeletal disorders that affect millions globally. The *Prdm12* gene encodes a conserved zinc finger transcriptional regulator expressed selectively in the nervous system. In humans, *PRDM12* mutations can cause congenital insensitivity to pain (CIP) or midface toddler excoriation syndrome (MiTES). Prdm12 is prominently expressed in developing somatosensory ganglia, where it plays a crucial role in nociceptive neuron development, its expression being maintained in mature C-LTMRs (C-low threshold mechanoreceptors) and nociceptive neurons. Despite enhanced understanding of Prdm12’s role in neuronal excitability and pain behavior, the impact of Prdm12 overexpression in mature nociceptive neurons has not been explored. Here, we conducted intravenous injection of AAV-PHP.S viral vectors encoding Prdm12-GFP (Prdm12-AAV) or GFP alone (Control-AAV), observing no overt changes in mouse behavior. When examining the properties of Prdm12 overexpressing sensory neurons in vitro, we observed an increase in rheobase alongside decreased neuronal responses to capsaicin and ATP, indicating a downregulation of TRPV1 and P2X ion channels activity, respectively. We next conducted intraarticular administration of viral constructs in female mice to determine how *Prdm12* overexpression in knee-innervating sensory neurons alters their excitability and influences inflammatory joint pain induced by intraarticular administration of complete Freund’s adjuvant (CFA). *Prdm12* overexpression in knee-innervating neurons decreased inflammation-induced changes in digging and weight bearing, prevented inflammation-induced neuronal hyperexcitability, and decreased macroscopic voltage-gated ion channel conductance. Our findings illustrate that Prdm12 overexpression strongly modulates neuronal excitability in adult animals, highlighting its importance in pain perception and its potential as an analgesic target.

**Summary:** Overexpression of the transcriptional regulator Prdm12 in knee-innervating neurons of mice reduces inflammatory joint pain and counteracts inflammation-induced neuronal hyperexcitability.

## 1 Introduction

The *Prdm12* gene encodes an evolutionarily conserved zinc finger transcriptional regulator selectively expressed in the nervous system. In humans, *PRDM12* mutations can result in either congenital insensitivity to pain (CIP) or midface toddler excoriation syndrome (MiTES)^1,2^. Prdm12 is highly expressed in developing somatosensory ganglia where it is selectively transcribed in all differentiating nociceptive neurons, specialized neurons that detect noxious stimuli, as well as C-low-threshold mechanosensitive neurons (C-LTMRs) that are activated in response to light mechanical stimulation and are involved in the sensation of touch^3,4^. Sensory neuron cell bodies are located in the dorsal root ganglia (DRG) and Prdm12 is required for DRG nociceptive neuron development due its role in the initiation and maintenance of the expression of the receptor for nerve growth factor (NGF), tropomyosin receptor kinase A (TrkA)^3,4^ and the repression of alternate somatosensory and visceral sensory determinants^5^. *Prdm12* remains expressed in mature C-LTMRs and nociceptive neurons where it contributes to the control of nociception^6,7^. *Prdm12* is also expressed in distinct regions and nuclei of the adult brain, including the arcuate nucleus of the hypothalamus where it is co-expressed with the pro-opiomelanocortin *Pomc* gene^8^. Despite recent research using knockout mice examining how Prdm12 regulates neuronal excitability and pain behaviour^6,7^, further investigation is required to better understand its function and mechanisms involved. Interestingly, complete Freund’s adjuvant (CFA)-induced knee inflammation causes decreased *Prdm12* expression in mature nociceptive neurons, and, in the absence of *Prdm12*, mice were hypersensitive to formalin-induced inflammatory paw pain^7^. Therefore, considering that decreased sensory neuron expression of *Prdm12* occurs during inflammatory pain and lack of sensory neuron Prdm12 is associated with hypersensitivity to certain noxious stimuli, we hypothesized that overexpression of *Prdm12* in sensory neurons might alleviate inflammatory pain.

Targeted AAV delivery via intra-articular injection provides a method for gene delivery to the nervous system to modulate joint pain locally^9–11^. The benefit of this approach is that it drives gene expression locally, thus mitigating against side effects associated with systemic drug administration. Therefore, here we used the AAV9 derivative AAV-PHP.S, which has been shown to display improved tropism towards peripheral neurons to selectively overexpress Prdm12 in transduced peripheral neurons^12^.

Our results reveal that intravenous AAV delivery of Prdm12 causes overexpression in DRG neurons, which resulted in no overt changes in behaviour, but did alter neuronal excitability. Furthermore, intraarticular AAV-mediated Prdm12 delivery to knee-innervating neurons prevented inflammation-induced hyperexcitability and pain behaviors. These results suggest a potent role of Prdm12 in regulating peripheral sensitization, thus accentuating its importance as a potential analgesic target.

## 2 Materials and Methods

### 2.1 Mice

C57BL/6J mice (Envigo) were provided ad libitum food and water and were housed at ∼21°C with a 12h light/dark cycle. For joint inflammation experiments, adult (8–12-week-old) female C57BL/6J mice were used because female sex increases the risk for arthritis^13^. The experimental protocols were regulated under the Animals (Scientific Procedures) Act 1986 Amendment Regulations 2012 following ethical review by the University of Cambridge Animal Welfare and Ethical Review Body, Project License P7EBFC1B1.

### 2.2 Intravenous and Intraarticular Knee injections

Two AAV-PHP.S viruses that use an AAV9 capsid for tissue targeting of sensory neurons and AAV2 inverted terminal repeats (ITRs) for genome packaging and a constitutive CAG promoter to overexpress the gene of interest were designed and produced by the viral vector facility at ETH Zurich: a Control-AAV, ssAAV-9/2-CAG-EGFP-WPRE-SV40p(A), (titer value = 2.9 x 10^13 vg/ml) that overexpresses EGFP alone (Fig. S1) and a Prdm12-AAV, ssAAV-9/2-CAG-EGFP-2A-3xFLAG-V5-mPrdm12-WPRE-SV40p(A) (titer value = 1.2 x 10^13 vg/ml) that overexpresses EGFP and a 3x Flag and V5 tagged version of Prdm12 (Fig. S2).

Intravenous injections of Control-AAV or Prdm12-AAV were administered via the lateral tail veins of conscious restrained animals (10^12 particles per injection, thus approximately 100 ul, depending on titer values).

Intra-articular injections were made through the patellar tendon with a 30G needle (10^11 particles per injection, thus approximately 10 ul, depending on titer values) bilaterally; mice were first anesthetized with ketamine (100 mg/kg) and xylazine (10 mg/kg), administered intra-peritoneally (IP). To induce joint inflammation, 10 μl CFA (10 mg/ml, Chondrex) was injected unilaterally into the knees of mice, 28 days after AAV injections. For this study there were 4 groups of mice: Control-AAV + CFA, Control-AAV + saline, Prdm-12-AAV + CFA, and Prdm-12-AAV + saline. To quantify inflammation, knee width was measured with digital Vernier’s calipers pre, and 24-hours post CFA injection.

### 2.3 Behavioral experiments

All experiments were carried out between 9:00-12:00. Before any behavioral experiments were conducted, mice were habituated in the procedure room in their home cages for 30 min. All experiments were conducted in presence of 2 researchers (1 female, 1 male), blinded to the conditions.

### 2.4 Digging

Digging behaviour was assessed as detailed previously^14^. In brief, mice were placed individually in standard 49 × 10 × 12 cm cages with a wire lid, filled with Aspen midi 8/20 wood chip bedding (LBS Biotechnology) tamped down to a depth of ∼4 cm containing Aspen midi 8/20 wood chip bedding (LBS Biotechnology). Mice were individually placed in the testing apparatus for a duration of 3 minutes without access to food and water to minimize distractions, and they were allowed to freely move and dig within the apparatus. Mice were video recorded for the entire duration of the experiment. The test was conducted twice; pre and 24-hours post CFA injection. The duration of digging was measured as the amount of time the mice actively displaced the bedding materials using their hind paws^15^. Additionally, the number of visible burrows at the end of the test period was recorded.

### 2.5 Dynamic weight bearing

Both humans^16^ and rodents^9,17^ exhibit changes in weight bearing during joint inflammation, reflecting pain-driven behavioral adaptations. Deficits in weight-bearing are indicative of spontaneous pain, and such deficits were assessed in freely moving mice using a dynamic weight bearing (DWB) device (BIOSEB, DWB2), which does not require animal restraint, thus significantly reducing stress levels that could impact experimental outcome. Mice were not previously trained on the device, being simply placed in the apparatus and recorded for 3 mins; a matrix comprising approximately 2000 high precision force sensors is embedded in the floor of the enclosure where the animal is free to move as it pleases. The system measures the weight distribution on all four paws. Paw prints from both forelimbs and hindlimbs were automatically identified using the two highest confidence levels of the device’s software, with at least 1 minute and 30 seconds of each 3-minute recording manually verified. Data analysis was performed by a single experimenter who was blinded to the experimental conditions.

### 2.6 Rotarod

The locomotor function and coordination of mice were assessed using a rotarod (Ugo Basile 7650). Mice underwent testing on a rotarod set at a constant speed of 7 rpm for 1 minute, followed by an accelerating program ranging from 7 to 40 rpm over a period of 5 minutes, totaling 6 minutes of testing. A similar protocol was employed for training the mice one day prior to the actual testing. During the testing sessions, mice were removed from the rotarod after two passive rotations or if they fell off. The test sessions were video recorded, and a blinded experimenter analyzed the videos to determine the latency (in seconds) to passive rotation or fall.

### 2.7 Hot/Cold plate

Mice were placed on the testing apparatus (BIOSEB), 20 cm diameter x 25 cm height, which consists of a temperature-controlled metal plate surrounded by a transparent acrylic cylinder. The metal plate was set at 55°C for hot sensitivity or 4°C for cold sensitivity. The latency to show a nociceptive response, e.g. hind paw lick, hind paw flick, or a jump was recorded, each mouse being immediately removed once a response was observed. If there was no response within 30 seconds, the test was terminated, and the mouse was removed from the hot/cold plate to prevent injury.

### 2.8 DRG neuron culture

Mice were euthanized by CO_2_ exposure followed by cervical dislocation. DRG were then isolated from dissected spinal columns in ice-cold L-15 Medium (1X) + GlutaMAX-1 (Life technologies) which was followed by incubation in 1 mg/ml type 1A collagenase for 15 minutes and then 1 mg/ml trypsin solution for 30 minutes at 37°C. DRG were then transferred to culture medium containing L-15 Medium (1X) + GlutaMAX-l, 10 % (v/v) fetal bovine serum, 24 mM NaHCO_3_, 38 mM glucose, 2 % (v/v) penicillin/streptomycin, and mechanically dissociated with a 1 ml Gilson pipette and briefly centrifuged. Supernatant containing dissociated DRG neurons was collected in a fresh tube. This cycle of mechanical trituration was repeated five times, after which the cells were plated on poly-D-lysine and laminin coated glass bottomed dishes (MatTek) and kept at 37°C, 5 % CO_2_ for 4-, 24-or 48-hours depending upon experimental need for electrophysiological recording and Ca^2+^-imaging. For experiments with mice following unilateral CFA knee injection, ipsilateral and contralateral DRG were kept separate throughout the dissociation. For immunocytochemistry experiments, dissociated DRG neurons were plated on poly-lysine/laminin coated glass coverslips (BD Biosciences) and incubated at 37 °C, 5% CO_2_.

### 2.9 Electrophysiology: Whole cell patch-clamp

All recordings were performed using a Multiclamp 700A amplifier and Digidata 1440A digitizer (Molecular devices). Patch pipettes of 2-5 MΩ were pulled with a P-97 Flaming/Brown puller (Sutter Instruments) from borosilicate glass capillaries. The extracellular solution (ECS) contained (in mM): NaCl (140), KCl (4), MgCl_2_ (1), CaCl_2_ (2), glucose (4) and HEPES (10), adjusted to pH 7.40 with NaOH, and an osmolality of 300-310 mOsm with sucrose. Patch pipettes were filled with intracellular solution containing (in mM): KCl (110), NaCl (10), MgCl_2_ (1), EGTA (1), HEPES (10), Na_2_ATP (2), Na_2_GTP (0.5), adjusted to pH 7.30 with KOH, and an osmolality of 310-315 mOsm with sucrose. Action potentials were generated by 40 ms current injections of –100-2000 pA in 21 pA steps. Rheobase, amplitude, half peak duration, afterhyperpolarization duration and amplitude were analyzed using Axon Clampfit software (Molecular Devices). The excitability of neurons was further assessed by applying a suprathreshold (2x action potential threshold) for 500 ms, the number of action potentials discharged during this time was counted.

The activity of voltage-sensitive channels was assessed in voltage clamp mode with appropriate compensation for series resistance. Cells were held at –120 mV for 100 ms before stepping to the test potential (–60 mV – 60 mV in 5 mV increments) for 40 ms and returning to a holding potential of –120 mV for 100 ms between steps to record tail currents. Peak inward and outward currents were normalized to cell size by dividing by cell capacitance.

Current-voltage (I-V) relationships were assessed by plotting peak current amplitudes against the corresponding command potentials. Conductance-voltage (G-V) curves were derived from the I-V data using the equation ΔG = ΔI/ΔV. I-V curves were fitted with a Boltzmann function to determine the reversal potential and half-activating potential of voltage sensitive channels.

### 2.10 Ca^2+^-imaging

For Ca^2+^-imaging analysis, we used the red-shifted Ca^2+^ indicator Rhod4 (ab112157 Abcam, Cambridge, UK) to avoid overlap with the AAV-derived GFP. DRG neurons were incubated with the dye for 30-mins at room temperature (RT). Neurons were then washed with ECS and placed on microscope for imaging using an inverted Nikon Eclipse Ti microscope, captured with a Zyla cSMOS camera (Andor) at 1 Hz, 50 ms exposure using Micro-Manager software (NIH). Agonists used were applied for 10-seconds in random order, with a 100s gap between the administration of either capsaicin (1 μm) or ATP (30 μm). KCl (50 mM) was used as a positive control to identify viable neurons.

For data analysis, KCl positive cells and one black background were drawn manually as regions of interest (ROI) using ImageJ software and the mean gray value of selected ROIs in sequence was extracted. Extracted data were then analyzed on R as previously described^17^. Briefly, the fluorescence signals of each cell were background and baseline corrected, only cells for which KCl elicited an increase in fluorescence > 5 x the SD of the baseline average were taken forward for further analyses. Fluorescence intensities were normalized to the maximal fluorescence elicited during KCl stimulation; cells were deemed capsaicin and/or ATP sensitive if fluorescence intensity surpassed the same threshold during the 10 second application of the agonist. Male mice were used in Ca^2+^ imaging experiments.

### 2.11 Immunohistochemistry

DRG were dissected in cold PBS, fixed for 20 minutes in 4% paraformaldehyde (PFA), then incubated overnight at 4°C in 30% sucrose. DRG were then mounted in Shandon M-1 Embedding Matrix (Thermo Fisher Scientific) and stored at –80°C until sectioning. Sections were cut at 14 μm using a Leica CM3000 cryostat. Sections were dried for 20 minutes at RT, rinsed 3 x 10 minutes with PBS-T (0.1% Triton X-100 in PBS), blocked with 10% normal donkey serum in PBS-T for 1 hour, and incubated with primary antibody in blocking solution overnight at 4°C. Primary antibodies used were: chicken anti-GFP 1:500 (Abcam, ab13970), and guinea pig anti-Prdm12 1:1000^4^. The next day, sections were rinsed 3 x 10 minutes with PBS and incubated with secondary antibodies diluted in blocking buffer at RT for 1 hour. Secondary antibodies used were donkey anti-chicken IgG Alexa Fluor 488 (1:1000) conjugated (Jackson laboratory, 703-545-155) and donkey anti-guinea pig IgG Alexa Fluor 594 (1:1000) conjugated (Molecular probes, A-11076). Sections were then rinsed 3 x 10 minutes in PBS-T, nuclei were stained with DAPI in PBS-T (1:2000) for 5 minutes, then washed 1 x 10 minutes with PBS-T, mounted with coverslips in Dako fluorescence mounting medium (DAKO cat. # S3023) and imaged using a wide-field fluorescence microscope Zeiss Axio Observer Z1 with the same parameters of acquisition per experiment. Image analysis was done using Fiji^18^. For Prdm12 fluorescence intensity analysis, ROIs of GFP+ve (that are DAPI+ve) neurons were used as cells of interest. Consistent thresholding parameters were used across all images to enable valid comparisons. After thresholding, binary masks were created and used to measure mean gray values. Mean grey values were then normalized to background values of cell-free regions.

### 2.12 Immunocytochemistry

Dissociated DRG neurons or HEK cells were plated onto 12 mm coverslips coated with poly-L-lysine and laminin. After 24 to 96 hours in culture, the cells were fixed at RT using 4% PFA for 10 minutes, followed by washing with PBS. Permeabilization of the cells was achieved by treating them with 0.05% Triton-X100 for 5 minutes at RT. Subsequently, cells were washed again with PBS and then incubated with a blocking buffer (1% donkey serum in 0.2% Triton-X100) for 30 minutes at RT. For immunostaining, cells were exposed to a primary antibody: chicken anti-GFP 1:500 (Abcam, ab13970), guinea pig anti-Prdm12 1:1000 and anti-Flag M2 1:500 (Cell Signaling, 8146), overnight at 4°C, following which they were washed with PBS and then incubated with secondary antibodies, diluted in PBS, with DAPI (1:1000), for 1 hour at RT. Secondary antibodies used were donkey anti-chicken IgG Alexa Fluor 488 (1:1000) conjugated (Jackson laboratory, 703-545-155), donkey anti-guinea pig IgG Alexa Fluor 594 (1:1000) conjugated (Molecular probes, A-11076), and donkey anti-mouse IgG Alexa Fluor 594 (1:600) conjugated (ThermoFisher, A-21203). Following a final wash, the coverslips were coverslips in Dako fluorescence mounting medium (DAKO cat#S3023) and imaged using a stereo microscope Olympus SZX16/Olympus BX51.

### 2.13 Statistical analysis

All statistical analyses were carried out using GraphPad Prism software (version 10) and R. Unless mentioned otherwise, values are represented as mean ± standard error of the mean (SEM). Statistical tests and number of mice used are specified in the legends of the figures wherever appropriate. Calculated p values are annotated.

## 3 Results

### 3.1 Prdm12 overexpression in sensory neurons does not influence animal behavior, but does influence neuronal chemosensitivity

We first tested the efficiency of *Prdm12* expression via AAV-PHP.S in HEK293 cells that normally do not express Prdm12^19^. Upon incubation with the viruses *in vitro*, ectopic Prdm12 expression was observed in cells transduced with Prdm12-AAV, but not those transduced with Control-AAV, whereas GFP expression was present following transduction with both viruses (Fig. S3). Proceeding to mice, Control-AAV and Prdm12-AAV were injected intravenously to induce GFP or *Prdm12* overexpression respectively in peripheral neurons of male and female mice (Figure 1A-B). Due to the observation that Prdm12 knockout induces a change in appetite^8^, body weight was measured, but no significant change occurred throughout the study (21 days) in any group (Fig. 1C). Neither locomotor performance (Fig. 1D) nor thermal nociception in response to either hot or cold stimuli (Fig. 1E-F) were affected by *Prdm12* overexpression, similar to previous findings in mice when *Prdm12* was ablated in DRG neurons^7^. We next assessed the function of DRG sensory neurons overexpressing *Prdm12*. However, we did observe differences between Control-AAV and Prdm12-AAV transduced neurons using Ca^2+^-imaging: DRG neurons from Prdm12-AAV injected mice showed a decreased likelihood to respond to capsaicin and ɑ,β-meATP in comparison to male mice injected with control-AAV (Fig. 1G-I).

**Figure 1:**
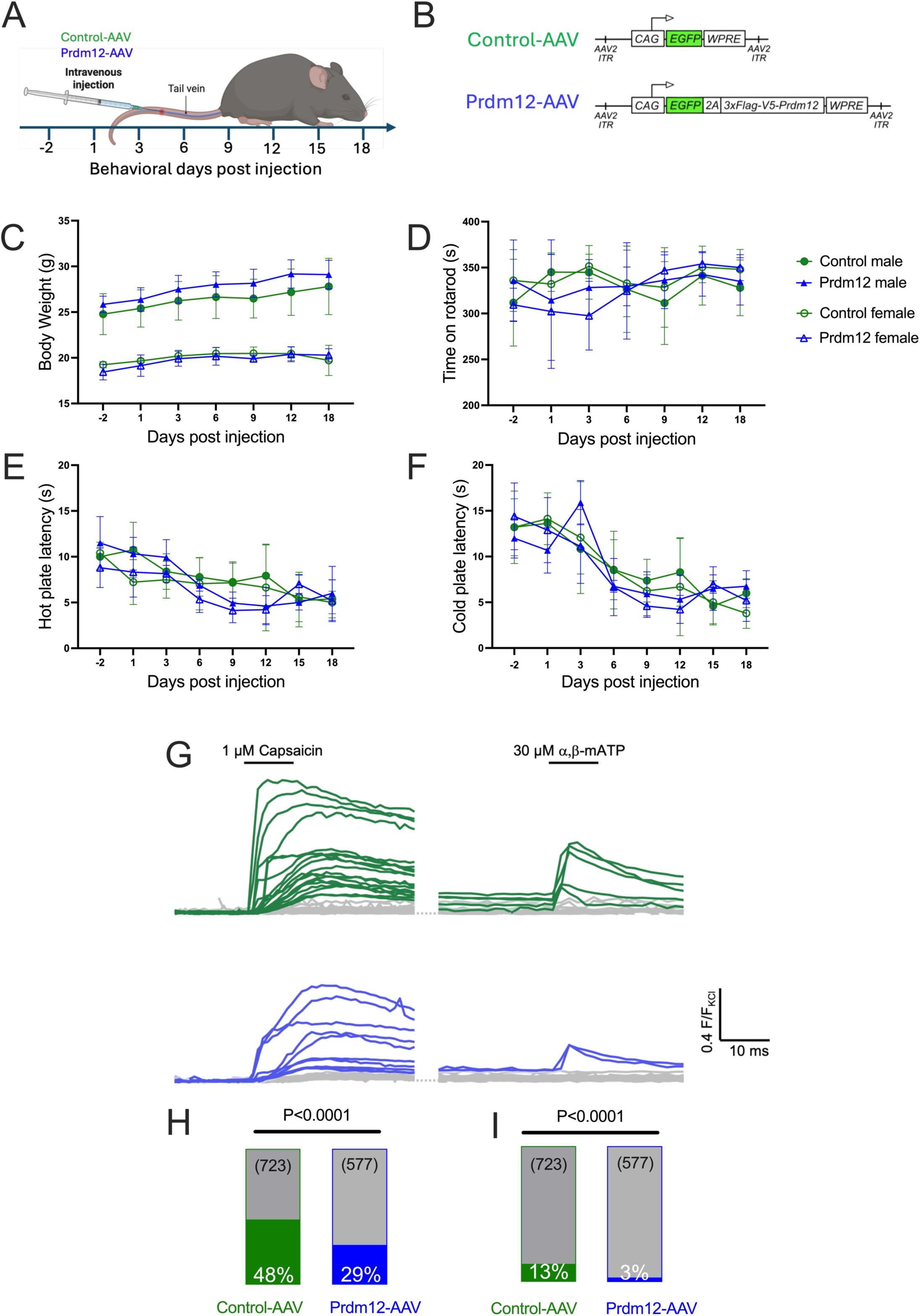
Prdm12 overexpression in peripheral neurons affects nociceptive neuron chemosensitivity without altering body weight, motor function or thermal sensitivity. (A) Schematic of the experimental design, illustrating the injection procedure. (B) Schematic of control-AAV and Prdm12-AAV constructs used for viral transduction. (C) Body weight measurements of male (closed symbols) and female (open symbols) mice injected with Control-AAV or Prdm12-AAV over 21 days (Control-AAV: male n=5, female n=5; Prdm12-AAV: male n=4, female n=5). (D) Rotarod performance (Control-AAV: male n=5, female n=5; Prdm12-AAV: male n=4, female n=5). (E) Hot plate response latencies at 55°C (Control-AAV: male n=5, female n=5; Prdm12-AAV: male n=4, female n=5). (F) Cold plate response latencies at 4°C (Control-AAV: male n=5, female n=5; Prdm12-AAV: male n=4, female n=5). (G) Representative Ca^2+^-imaging traces from cultured DRG neurons of Control-AAV (top) and Prdm12-AAV (bottom) injected mice in response to capsaicin (1 µM) and ATP (30 µM). Traces of 40 random cells were selected, positive responders are coloured, non-responders are in grey. (H) Percentage of cultured DRG neurons responding to capsaicin and (I) ATP from Control-AAV and Prdm12-AAV injected male mice (Control-AAV: 577 neurons from 4 mice; Prdm12-AAV: 723 neurons from 4 mice). Chi-square test.

### 3.2 Prdm12 overexpression in sensory neurons increases rheobase

Neuronal excitability can be influenced by duration of culture time and culture medium constituents^20–24^, therefore all subsequent experiments with cultured dissociated DRG neurons were conducted between 12-24-hours following initial plating.

AAV transduction efficiency was first tested in vitro. Lumbar DRG were dissociated and neurons cultured in the presence of Control-AAV or Prdm12-AAV viruses (1 x 10^11 vg/ml, Fig. 2A). 24-hours later, immunocytochemistry experiments revealed a robust signal for Prdm12 and GFP in neurons cultured with both viruses; Prdm12 is naturally expressed in all nociceptive neurons and C-LTMRs, i.e. expression is expected even in Control-AAV cultures (Fig. 2B). Subsequently, GFP-expressing neurons were recorded using whole cell patch clamp electrophysiology, current clamp recordings first being used to study the voltage changes produced by ion channel activity in the cell membrane that underpin action potential (AP) electrogenesis^25,26^. Current clamp tests showed a small, but significant, increase in rheobase, the minimum current that elicits an AP, in neurons overexpressing Prdm12 vs. control neurons, although there was no change in the resting membrane potential (Fig. 2C-D) and, other than rheobase, no AP parameter was significantly altered (Table 1). A higher rheobase reflects the need for a greater current injected to elicit an action potential and reflects a decrease in neuronal excitability. Thus, Prdm12 overexpression appears to cause a decrease in sensory neuron excitability. We examined this further by measuring macroscopic voltage-gated currents. GFP-expressing neurons transduced with Prdm12-AAV exhibited significantly smaller peak amplitude macroscope voltage-gated inwards current (Fig. 2E-F) and significantly smaller peak amplitude macroscope voltage-gated outward currents (Fig. 2G-H).

**Figure 2:**
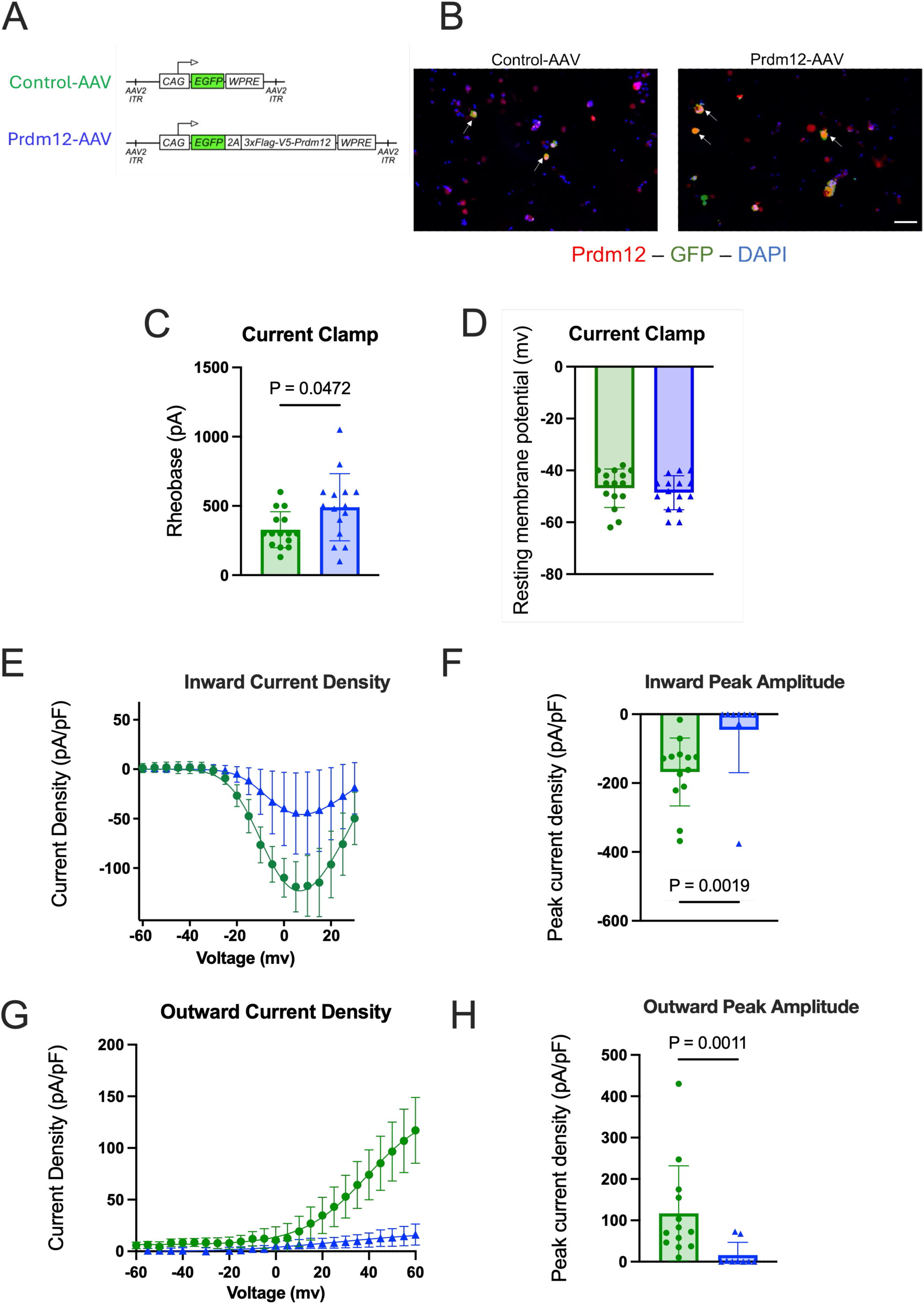
*Prdm12* overexpression *in vitro* increases rheobase of dissociated DRG neurons isolated from female mice. (A) Schematic of Control-AAV and Prdm12-AAV constructs used for viral transduction. (B) Representative images of cultured dissociated DRG neurons 24 hours following AAV transduction. GFP (green), Prdm12 (red), and DAPI (blue) labels nuclei. Arrows point to GFP+ve Prdm12 expressing cells. Scale bar: 100 μm. (C) Rheobase measurements in Control vs Prdm12-overexpressing neurons 12-24 hours after culture. Data are presented as mean ± SEM, Mann-Whitney test. (D) Resting membrane potential measurements in Control vs Prdm12-overexpressing neurons 12-24 hours after culture. Data are presented as mean ± SEM, Mann-Whitney test. (E) Current density for voltage-gated inward currents (pA/pF). (F) Voltage-gated inward current peak amplitude (pA/pF). (G) Current density for voltage-gated outward currents (pA/pF). (H) Voltage-gated outward current peak amplitude (pA/pF)

**Table 1:**
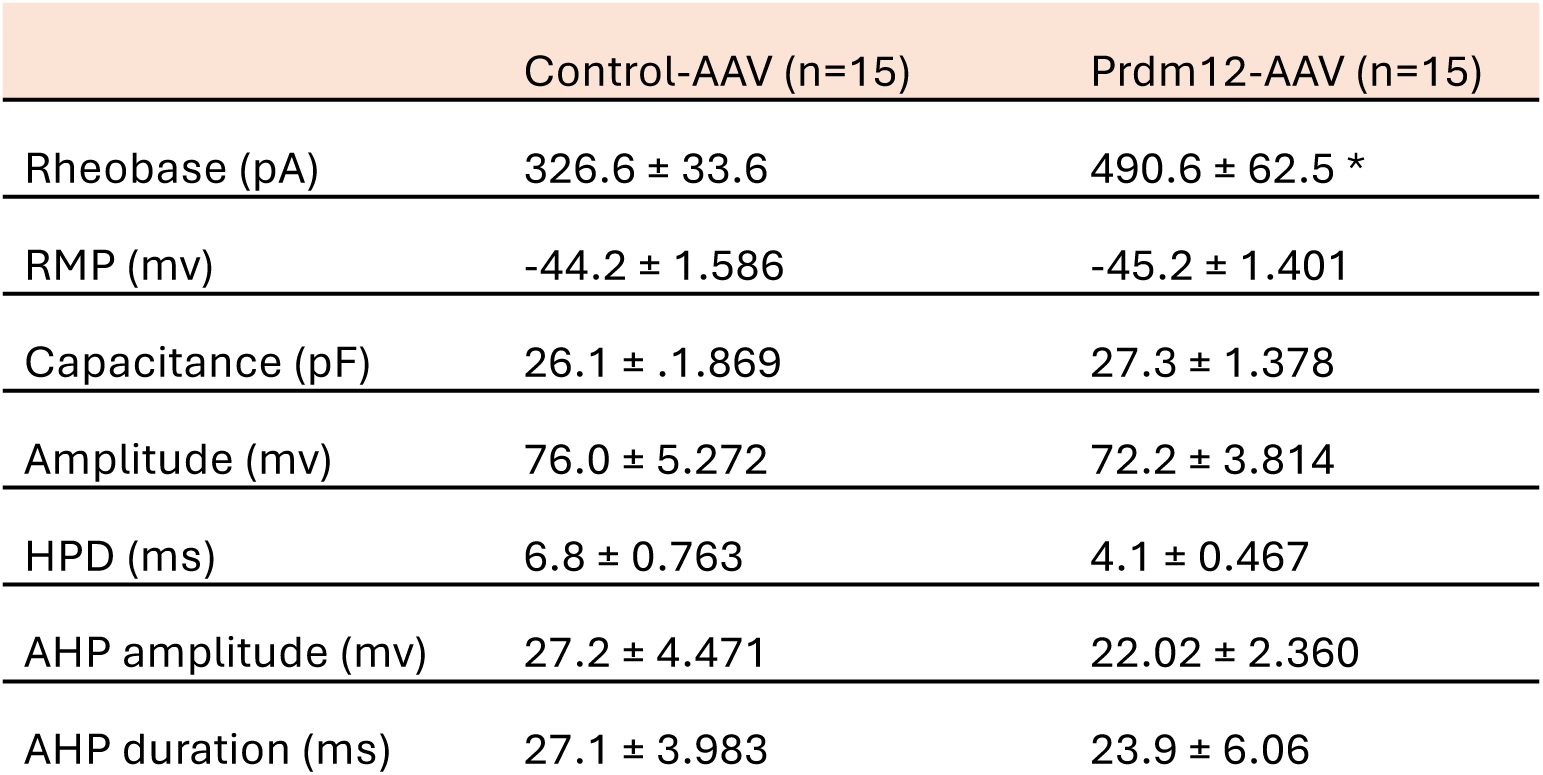
Action potential properties of DRG neurons transduced *in vitro* with AAV-PHP.S. Values are mean ± SEM, Mann-Whitney test * = in comparison to control-AAV neurons RMP: resting membrane potential; HPD: half peak duration; AHP: afterhyperpolarization *p=0.0472

### 3.3 Transduction efficiency of lumbar DRG neurons by knee-injected AAV-PHP.S

Having observed that Prdm12-AAV transduces DRG neurons in vitro and causes an increase in rheobase, we hypothesized that this increase in rheobase might counteract neuronal sensitization that occurs during inflammation in vivo. Therefore, we first injected either Control-AAV or Prdm12-AAV into both knees of mice (Fig. 3A-B) and observed a similar proportion of GFP labelling in the lumbar (L2-L5) DRG neurons (n = 4, all female) for mice in Control-AAV or Prdm12-AAV groups (Fig. 3C-D). Prdm12 is endogenously expressed in all nociceptive neuron subtypes as well as C-LTMRs, i.e. it is expressed in the majority of DRG sensory neurons^27^. Although the number of GFP+ve neurons co-expressing Prdm12 appears similar in Prdm12-AAV vs Control-AAV injected mice (Fig. 3E), the Prdm12 intensity was increased by ∼1.5 fold in GFP+ve neurons infected with Prdm12-AAV compared to those infected with Control-AAV, thus demonstrating an increase in Prdm12 expression following Prdm12-AAV administration (Fig. 3F).

**Figure 3:**
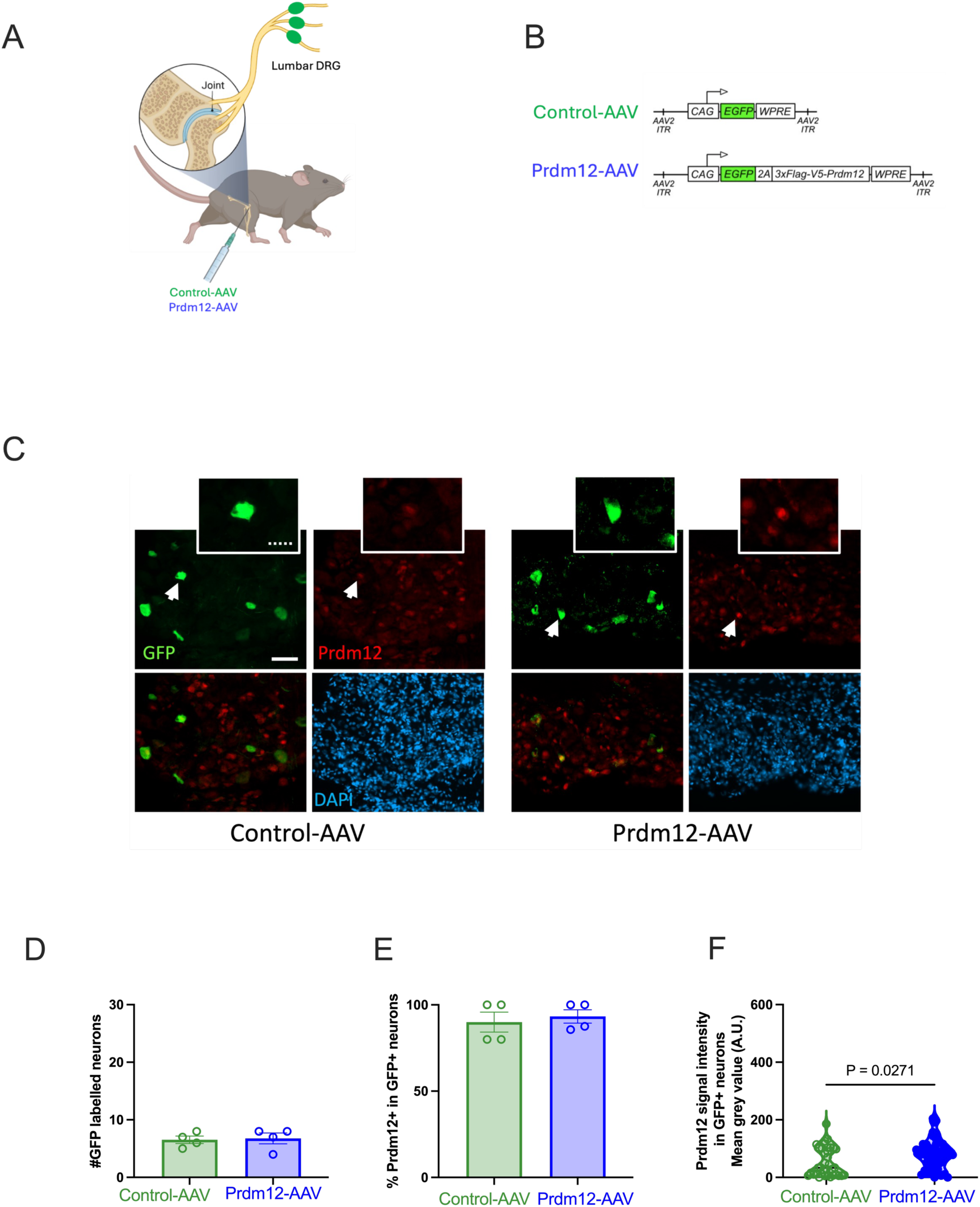
Transduction efficiency of DRG neurons following intra-articular knee injection of Control-AAV or Prdm12-AAV in female mice. (A) Schematic of the experimental design, illustrating the injection procedure. (B) Overview of the AAV-PHP.S viruses used in the study: Control-AAV, which induces GFP expression, and Prdm12-AAV, which induces both GFP and Prdm12 expression. (C) Immunostaining of sections of lumbar DRG showing the expression of GFP (green) and Prdm12 (red) in traced neurons. Scale bar: 50 μm. Dashed scale bar in inset: 25 μm. (D) Quantification of the number of GFP+ve neurons in the L2–L5 DRG labeled with AAV-PHP.S. (E) Percentage of Prdm12+ve neurons among the GFP+ve neurons. (F) Comparison of Prdm12 intensity between GFP+ve neurons in mice injected with Control-AAV or Prdm12-AAV. Bars represent mean ± SEM from 4 female mice. Statistical significance was assessed using an unpaired t-test.

### 3.4 CFA-induced knee inflammation

Having established that Control-AAV or Prdm12-AAV efficiently transduced knee-innervating neurons, we injected adult female mice with either Control-AAV or Prdm12-AAV intra-articularly in both knee joints. 4-weeks post-virus administration, half the mice in each group received a unilateral CFA knee injection and the other half received saline; the knee receiving CFA/saline is referred to as the ipsilateral knee and the other, non-injected knee the contralateral (Fig. 4A-B). To assess the effects of Prdm12 overexpression on joint pain, behavioral measures were made pre– and post-CFA/saline injection. Robust swelling was observed 24 hours post-CFA injection in the ipsilateral knee, while the contralateral knee showed no swelling, no inflammation being observed following saline injection, and no difference was observed for the knee width of animals injected with either Control-AAV or Prdm12-AAV prior to saline/CFA injection, i.e. AAV injections did not induce any inflammation and CFA-induced inflammation was of a similar magnitude in both Control-AAV and Prdm12-AAV injected mice (Fig. S4). Spontaneous pain, a common phenotype of arthritic disorders, significantly diminishes the overall feeling of well-being, which can be reflected by changes in natural pain behavior and well-being in rodents. Digging behavior and dynamic weight bearing (DWB) are both valuable experimental techniques to measure spontaneous pain in a stress-free non-restrained manner. For example, CFA-induced knee inflammation decreases mouse digging behavior and the hind paw weight placed on the CFA-injected limb^9,11,15,22^. Control-AAV mice receiving CFA showed a decrease in digging time as well as a significant decrease in the number of burrows dug, but no such change occurred in mice receiving Prdm12-AAV (Fig. 4C-D). As has been shown previously^9,28^, unilateral CFA injection did not alter rotarod performance in any group in this study, reflecting no gross change in motor coordination (Fig. 4E).

**Figure 4:**
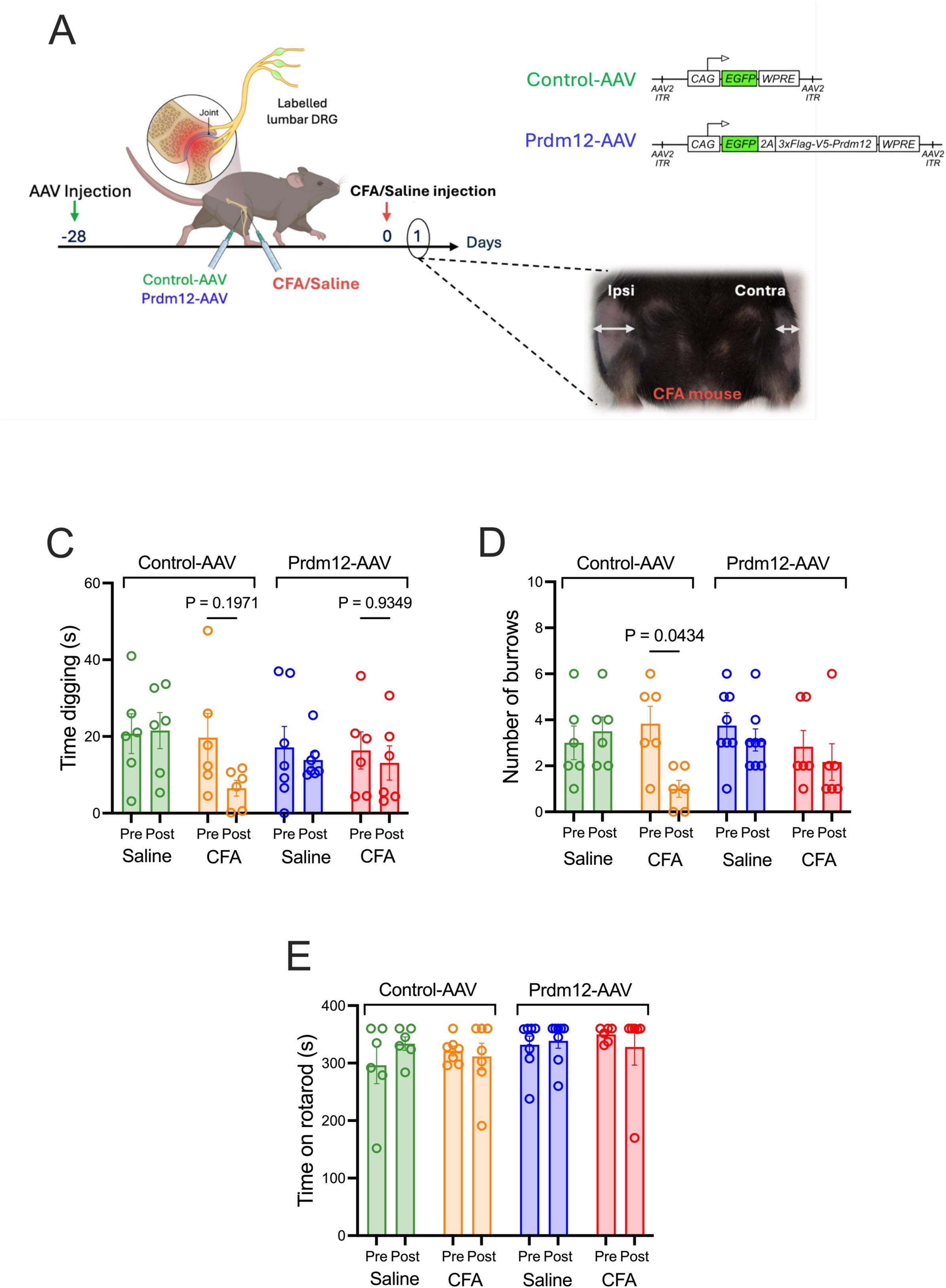
Assessment of motor coordination and digging behavior of female adult mice following intra-articular knee injection with Control-AAV or Prdm12-AAV and subsequent CFA or Saline. (A) Experimental timeline with a representative image of a mouse injected firstly with Control-AAV or Prdm12-AAV, followed by subsequent CFA or saline injection on the ipsilateral side (Ipsi) side. Arrows indicate the measurement points for knee width. (B) Overview of the AAV-PHP.S viruses used in the study: Control-AAV, which induces GFP expression, and Prdm12-AAV, which induces both GFP and Prdm12 expression. (C) Digging time (seconds) and (D) number of burrows, reflecting natural digging behavior, pre– and 24 hours post-CFA or saline injection in Control-AAV and Prdm12-AAV injected mice. (E) Time spent on the rotarod, reflecting motor coordination, pre– and 24 hours post-CFA or saline injection in mice injected with Control-AAV or Prdm12-AAV. Sample sizes: Control-AAV + Saline n=6; Control-AAV + CFA n=7; Prdm12-AAV + Saline n=8; Prdm12-AAV + CFA n=6. Data were collected over two independent experiments of AAV injections. Definitions: Contra = contralateral; Ipsi = ipsilateral. Data are presented as mean ± SEM. Statistical analysis was performed using a two-way ANOVA followed by Bonferroni post-hoc test.

When examining weight bearing, Control-AAV injected animals receiving CFA exhibited a shift in hind paw weight bearing such that significantly less weight was placed on the ipsilateral hind paw, which was not observed in saline injected mice (Fig. 5A-B). By contrast, no such significant decrease in ipsilateral hind paw weight bearing was observed following CFA injection in Prdm12-AAV mice and nor was any change observed following saline injection (Fig. 5C-D). We also examined fore paw weight bearing, but no changes were observed in CFA or saline injected mice in either Control-AAV or Prdm12-AAV groups (Fig. 5E-H), results indicating that changes in hind paw weight bearing were due to the CFA-induced inflammation. Further examination revealed that CFA injection of Control-AAV injected mice led to a decrease in ipsilateral/contralateral weight of hind paws as well as the ipsilateral/contralateral occupied area, neither of which occurred in Prdm12-AAV injected mice (Fig S5A and B). No change was observed regarding fore/hind paw weight or fore paws occupied area, similarly, rearing weight and duration, when mice are standing on both hind paws in an upright position, was unaffected in all conditions (Fig S5 C-F). These results suggest that Prdm12 overexpression could have role in regulating CFA-induced pain-related behaviors.

**Figure 5:**
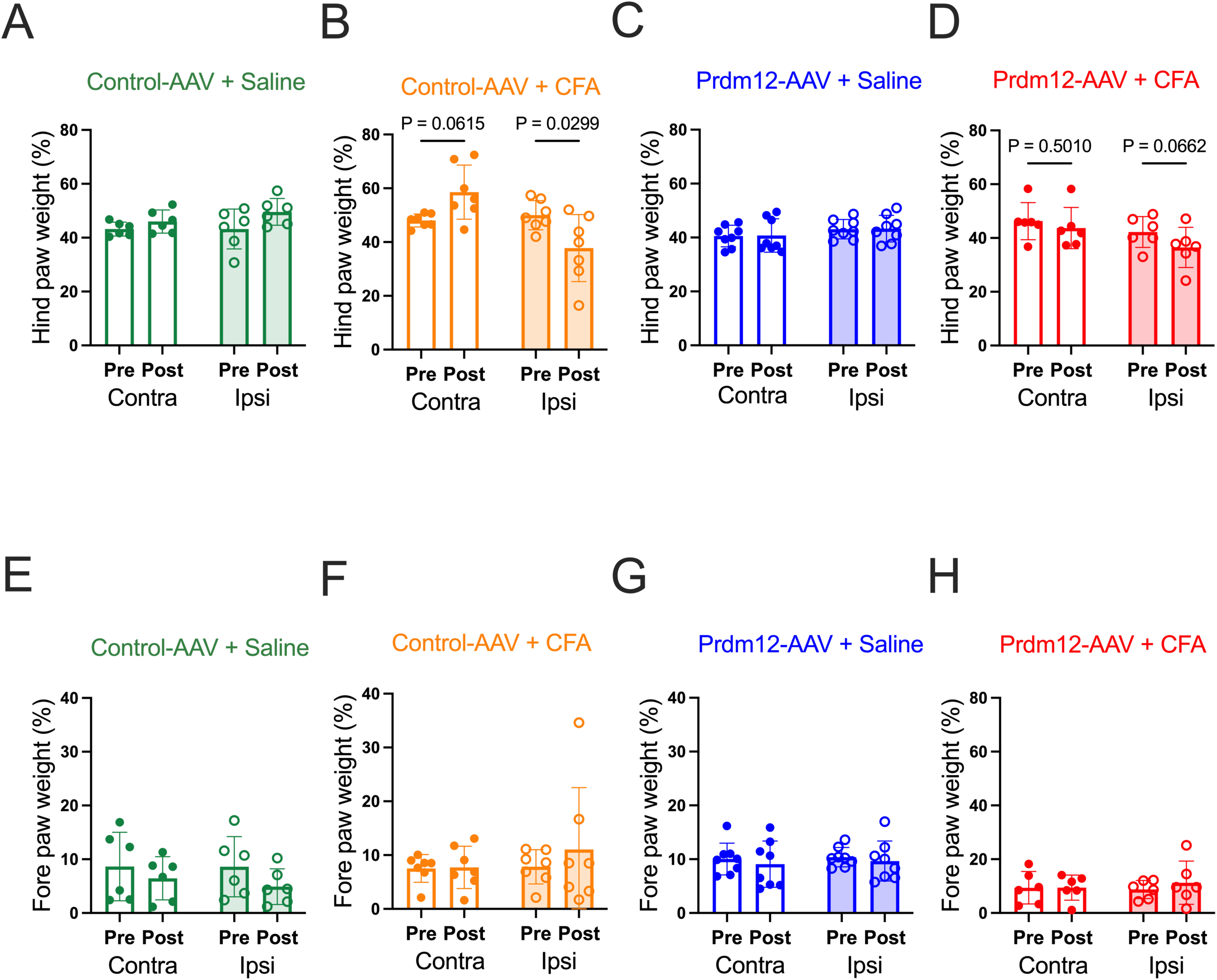
Dynamic weight bearing comparison of female adult mice following intra-articular knee injection with Control-AAV or Prdm12-AAV and subsequent CFA or Saline. (A-D) Percentage of hind paw weight-bearing, pre– and 24 hours post-CFA or saline injection in mice injected with Control-AAV or Prdm12-AAV. (E-H) Percentage of forepaw weight-bearing, pre– and 24 hours post-CFA or saline injection in mice injected with Control-AAV or Prdm12-AAV. Sample sizes: Control-AAV + Saline n=6; Control-AAV + CFA n=7; Prdm12-AAV + Saline n=8; Prdm12-AAV + CFA n=6. Data were collected over two independent experiments of AAV injections. Definitions: Pre = prior to CFA/saline injections; Post = 24 hours following CFA/saline injections; Contra = contralateral; Ipsi = ipsilateral. Data are presented as mean ± SEM. Statistical analysis was performed using a two-way ANOVA followed by Bonferroni post-hoc test.

### 3.5 Prdm12 overexpression decreases knee-afferent neuronal excitability in inflammatory conditions

Intraarticular administration of Prdm12-AAV prevented CFA-induced pain-related behaviors and CFA is known to induce knee-innervating neuron hyperexcitability^22^. We therefore hypothesized that Prdm12-AAV induction of Prdm12 expression might prevent CFA-induced hyperexcitability of ipsilateral knee-innervating neurons and thus explain the differences in inflammation-induced behaviors between Control-AAV and Prdm12-AAV groups; DRG neurons innervating the contralateral knee were used as control. We recorded electrically evoked APs from neurons with similar sizes, as indicated by comparable cell capacitances (Table 2), measuring rheobase (Fig. 6A-B). Ipsilateral knee-innervating neurons isolated from mice injected with Control-AAV had a more depolarized resting membrane potential than knee-innervating contralateral neurons (Fig. 6C). By contrast, the CFA-induced depolarization of the resting membrane potential was not observed in ipsilateral knee-innervating neurons isolated from mice injected with Prdm12-AAV (Fig. 6C); no difference was observed between resting membrane potentials of neurons innervating the contralateral side of Control-AAV vs. Prdm12-AAV injected mice, i.e. as we observed following in vitro Prdm12-AAV transduction (Table 1), Prdm12 overexpression does not appear to alter the resting membrane potential under healthy conditions, but rather counteracts the inflammation-induced decrease (Fig. 6C). Similarly, the rheobase of Control-AAV neurons innervating the CFA-injected knee (Ipsi) was lower than that of Control-AAV neurons innervating the non-injected knee (Contra), a finding in line with our previous observations^9,22^, indicative of neuronal sensitization (Fig. 6B). However, as was observed with Prdm12 overexpression in vitro (Fig. 2C), Prdm12-AAV neurons innervating the contralateral/non-injected knee had a higher rheobase than Control-AAV neurons innervating the contralateral/non-injected knee, and, although a decrease in rheobase was observed in Prdm12-AAV neurons innervating the ipsilateral/CFA-injected knee vs. Prdm12-AAV neurons innervating the contralateral/non-injected knee, this was not significant, the rheobase being similar to that measured in Control-AAV neurons innervating the contralateral/non-injected knee (Fig. 6B). One difference observed in Prdm12-AAV neurons that was not observed in previous *in vitro* experiments was a significant increase in AP amplitude in contralateral and ipsilateral neurons compared to Control-AAV equivalents (Fig. 6D). No other AP parameters were altered (Table 2). Lastly, when challenged with a suprathreshold stimulus, Control-AAV contralateral neurons exhibited a greater number of APs than Prdm12-AAV contralateral neurons, again suggestive of Prdm12 overexpression counteracting inflammation-induced neuronal hyperexcitability (Fig. 6E-F).

**Figure 6:**
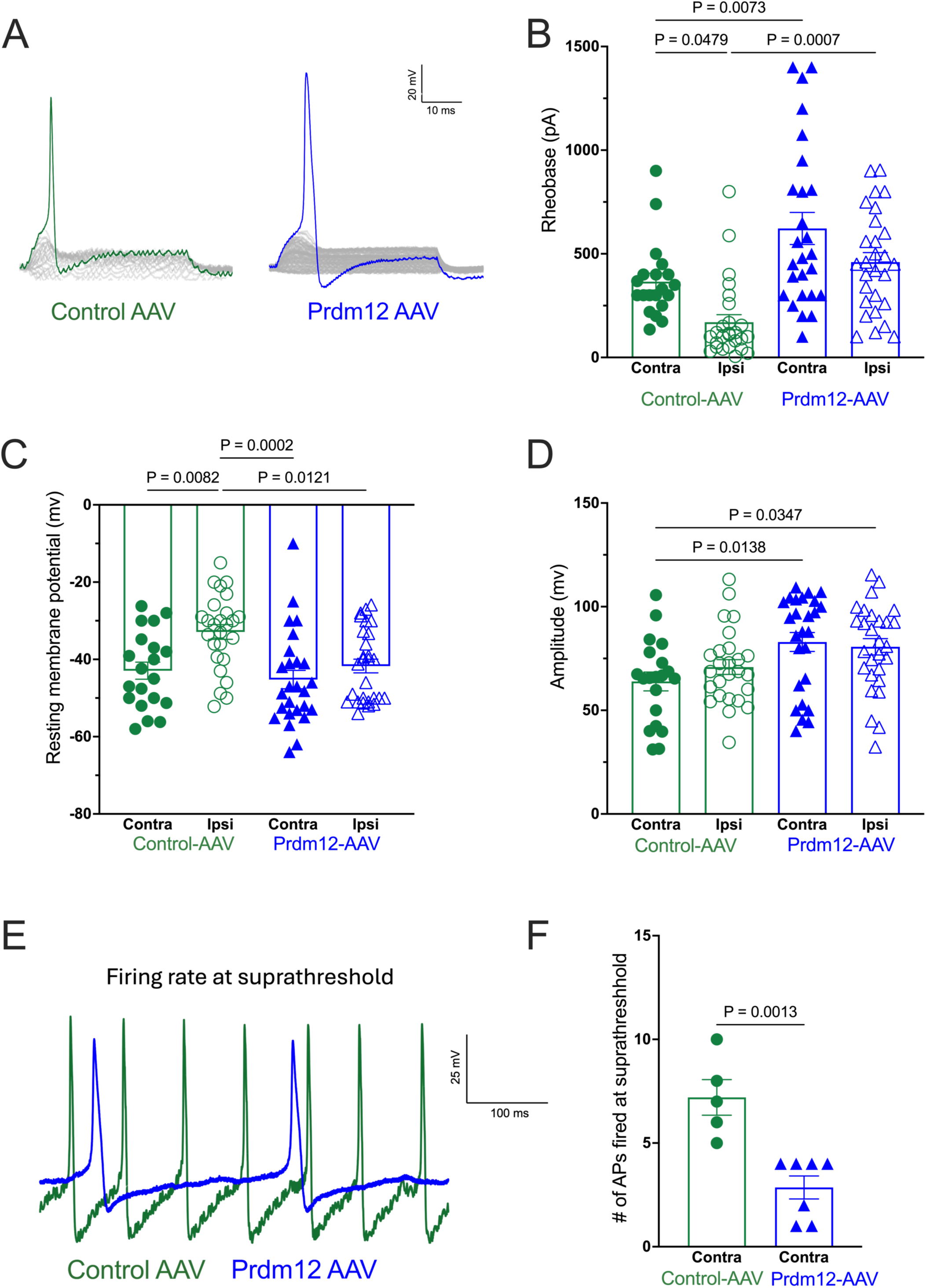
Prdm12 overexpression prevents CFA-induced neuronal hyperexcitability of knee-innervating afferent DRG neurons Analysis of action potential properties of knee-innervating DRG neurons from female mice injected with Control-AAV or Prdm12-AAV in both knees, and subsequent unilateral CFA injection, comparing neurons from the CFA-injected side (Ipsi) with those innervating the non-injected side (Contra): (A) Representative action potential traces in response to current injection, (B) Rheobase (pA), (C) Resting membrane potential (mV), (D) Action potential amplitude (mV), (E) Representative suprathreshold (2x rheobase) action potential traces, and (F) Number of action potentials generated in neurons stimulated at suprathreshold stimulation (2x rheobase). Data were collected from 4 female mice per group over 3 independent experiments involving AAV injections followed by DRG neuronal dissociation. Sample sizes: Control-AAV n=4; Prdm12-AAV n=5. Statistical analysis: One-way ANOVA followed by Tukey’s multiple comparison test. Data are presented as mean ± SEM. Abbreviations: Ipsi = Ipsilateral (CFA-injected side); Contra = Contralateral (non-injected side)

**Table 2:**
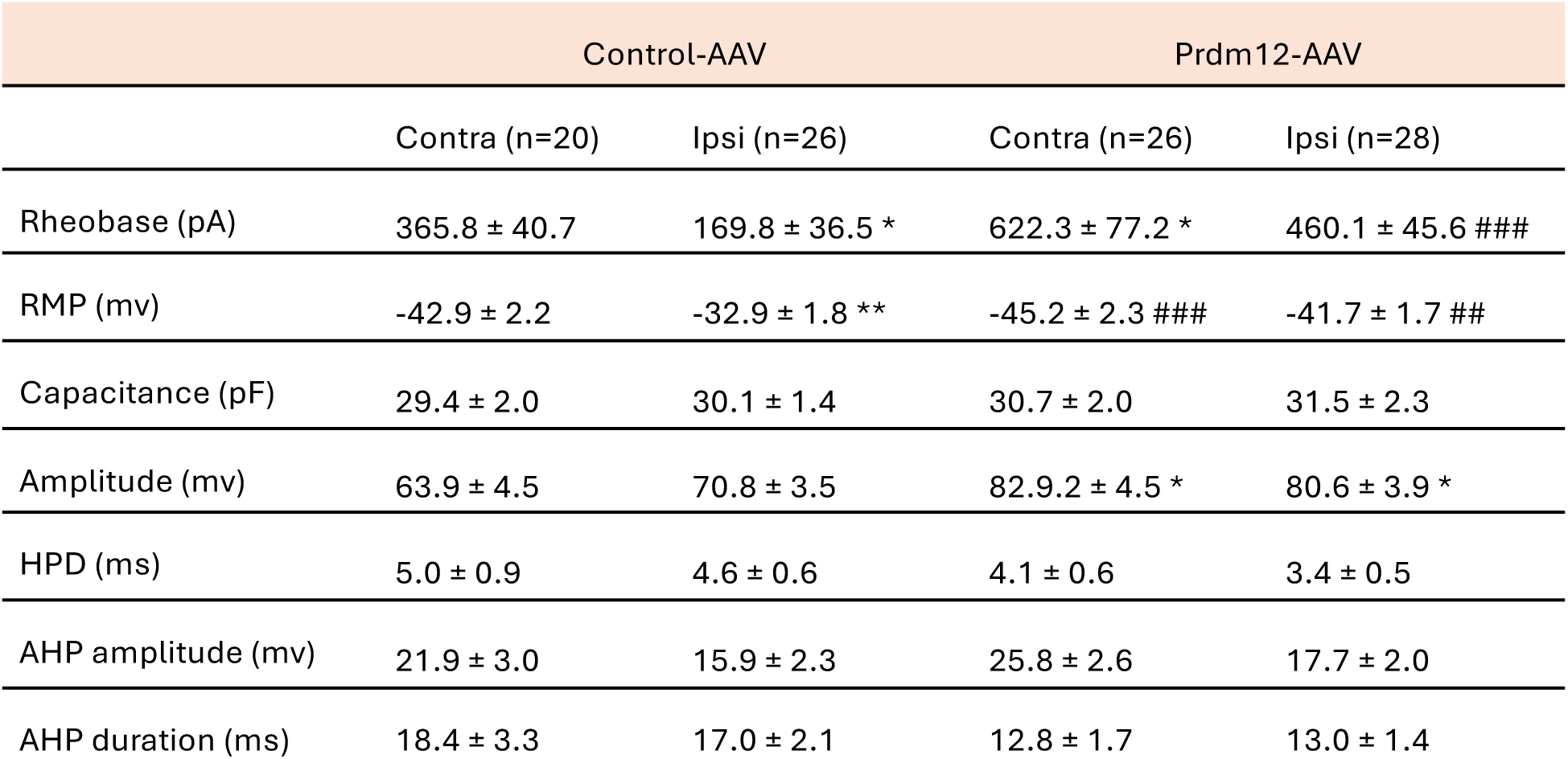
Action potential properties of knee-innervating DRG neurons following intra-articular injection with AAV-PHP.S constructs and subsequent CFA injection. 4 female mice were used per condition. Values are mean ± SEM, Anova test followed by Tukey’s multiple comparison test * = in comparison to control-AAV neurons injected with Saline (Control-AAV/ Contra) # = in comparison to control-AAV neurons injected with CFA (Control-AAV/ Ipsi) */# indicates p < 0.05, **/## indicates p < 0.01, ***/### indicates p < 0.0001. Ipsi: Ipsilateral; Contra: Contralateral; RMP: resting membrane potential; HPD: half peak duration; AHP: afterhyperpolarization

|  | Control-AAV |  | Prdm12-AAV |  |
| --- | --- | --- | --- | --- |
|  | Contra (n=20) | Ipsi (n=26) | Contra (n=26) | Ipsi (n=28) |
| Rheobase (pA) | 365.8 ± 40.7 | 169.8 ± 36.5 * | 622.3 ± 77.2 * | 460.1 ± 45.6 ### |
| RMP (mv) | -42.9 ± 2.2 | -32.9 ± 1.8 ** | -45.2 ± 2.3 ### | -41.7 ± 1.7 ## |
| Capacitance (pF) | 29.4 ± 2.0 | 30.1 ± 1.4 | 30.7 ± 2.0 | 31.5 ± 2.3 |
| Amplitude (mv) | 63.9 ± 4.5 | 70.8 ± 3.5 | 82.9.2 ± 4.5 * | 80.6 ± 3.9 * |
| HPD (ms) | 5.0 ± 0.9 | 4.6 ± 0.6 | 4.1 ± 0.6 | 3.4 ± 0.5 |
| AHP amplitude (mv) | 21.9 ± 3.0 | 15.9 ± 2.3 | 25.8 ± 2.6 | 17.7 ± 2.0 |
| AHP duration (ms) | 18.4 ± 3.3 | 17.0 ± 2.1 | 12.8 ± 1.7 | 13.0 ± 1.4 |

As for in vitro transduction experiments, we also examined how *Prdm12* overexpression affected voltage-gated currents, observing a significant reduction of both inward and outward macroscopic voltage-gated current densities in contralateral neurons isolated from Prdm12-AAV mice vs. those isolated from Control-AAV mice. (Fig. 7A-E). However, closer examination demonstrated that Prdm12 overexpression significantly reduced only macroscopic voltage-gated inward conductance, where a rightward shift within the activation range of inward ion channels indicates a change in the voltage dependence of channel activation, i.e. requiring a more positive membrane potential to open. This decrease in inward conductance likely contributes to the decreased neuronal excitability of cells overexpressing Prdm12 (Fig. 7D-G).

**Figure 7:**
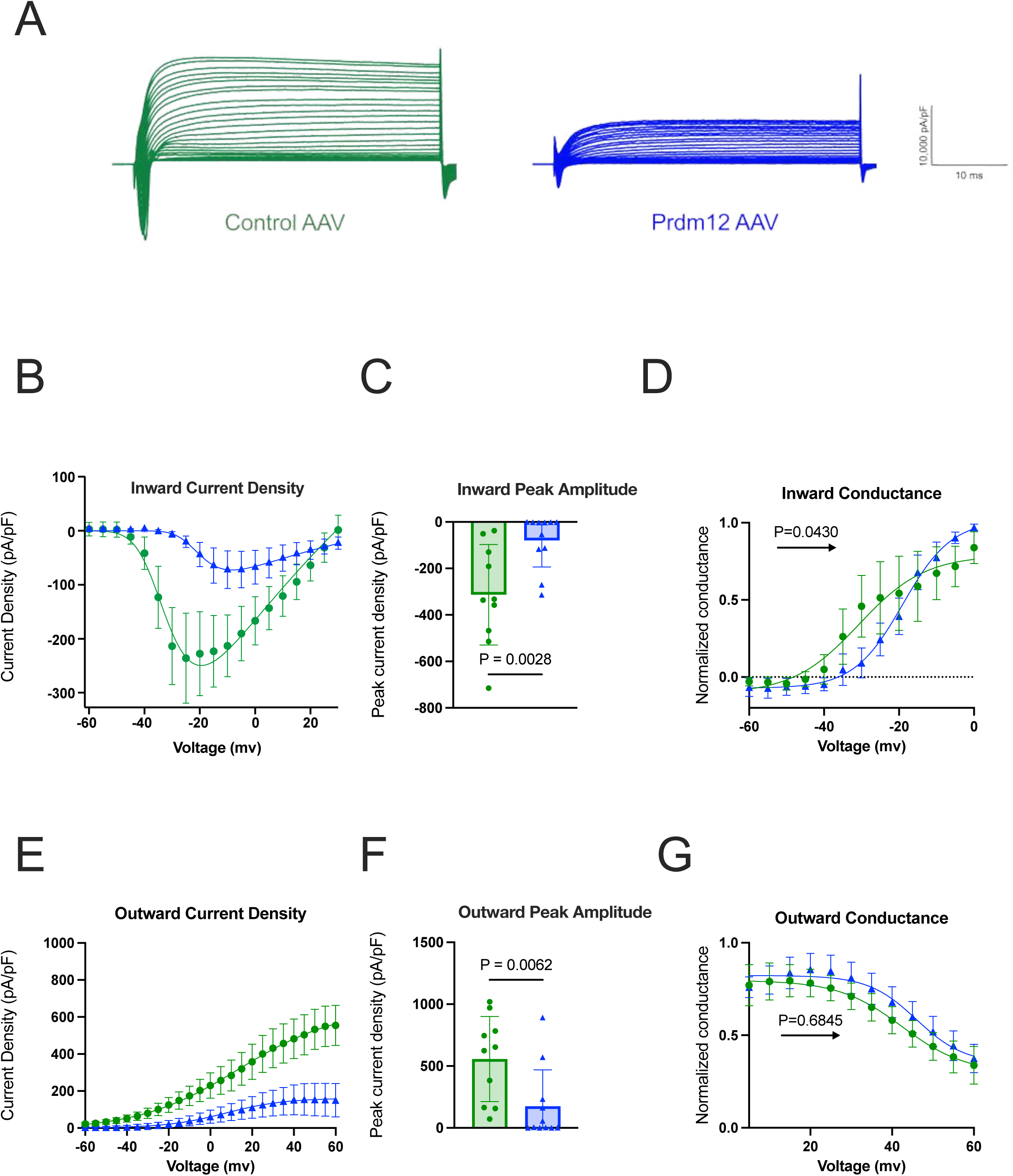
Prdm12 overexpression modulates voltage-gated currents in knee-innervating DRG neurons Analysis of inward and outward current properties of contralateral (non-injected side) knee-innervating DRG neurons from female mice injected with Control-AAV or Prdm12-AAV: (A) Representative I-V current traces from control-AAV and Prdm12-AAV transduced neurons, (B) Current density for voltage-gated inward currents, (C) Voltage-gated inward current peak amplitude (pA/pF), (D) Normalized conductance for voltage-gated inward currents, (E) Current density for voltage-gated outward currents (pA/pF), (F) Voltage-gated outward current peak amplitude (pA/pF), and (G) Normalized conductance for voltage-gated outward currents. Data are presented as mean ± SEM. Statistical analysis was performed using unpaired t-test n = 10 neurons from Control-AAV and n = 11 neurons from Prdm12-AAV. *p < 0.05, **p < 0.01, ***p < 0.001.

## 4 Discussion

Mutations in Prdm12 can lead to CIP and MiTES in humans^1,2^, thus demonstrating its key role in somatosensation. Moreover, unlike other painlessness genes, the expression of Prdm12 is almost exclusive to nociceptive neurons in adults, which makes it an attractive therapeutic target^29,30^. Constitutive Prdm12 knockout is lethal, but Prdm12 inducible conditional knockout (iCKO) mouse models have shown that Prdm12 is not required for cell survival in adulthood but continues to play a role in the transcriptional control of a network of genes, many encoding ion channels and receptors^6,7^. Furthermore, hypothalamic Prdm12 has been shown to regulate Pomc expression, as well as food intake, adiposity, and weight^8^. Nonetheless, ablating Prdm12 expression in adults had no effect on locomotion, anxiety, or hippocampal-dependent learning^6^.

Despite the extensive body of research elucidating the effects of *Prdm12* ablation in DRG neuron development and function, the effect of *Prdm12* overexpression on mature nociceptive neurons has not yet been explored. Here we show, firstly via intravenous Prdm12-AAV administration, that overexpression of *Prdm12* in peripheral neurons of male mice causes a significant decrease in neuronal response to algogens: capsaicin and ATP. Secondly, inducing Prdm12 overexpression in knee-innervating afferents causes a decrease in inflammation-induced changes in digging and weight bearing, concomitant with an absence of inflammation-induced sensitization of knee-innervating neurons in female mice. These results align with a previous study where CFA-induced inflammation decreased *Prdm12* expression in mature nociceptive neurons, and where it was also shown that in the absence of *Prdm12* mice were hypersensitive to formalin-induced inflammatory pain^7^. Dysregulated expression of *Prdm12* has also been observed in response to nerve injury^31^, and in pruritoceptive neurons in response to dermatitis^32^. Moreover, mice lacking *Prdm12* exhibited a reduced response to capsaicin, which activates TRPV1 on nociceptive C-fibers, generating pain^6,7^. However, in the absence of Prdm12, normal responses were observed in response to noxious thermal and mechanical stimuli^6,7^, which is consistent with our observation of a lack of altered thermal sensitivity when *Prdm12* was overexpressed in peripheral neurons. Thus, Prdm12 may not play a critical role in modulating baseline thermal nociception or the development of thermal hyperalgesia in these contexts. Furthermore, a comparative transcriptome profiling study of human DRG in pain conditions has shown that *Prdm12* expression generally decreases in pain conditions in females and increases in males suggesting sexual dimorphism in Prdm12 role in pain, which means sexual grouping should be accounted for in studies exploring Prdm12 role, especially in relation to pain^23^.

When transducing DRG sensory neurons *in vivo*, *Prdm12* overexpression drove an increase in rheobase, i.e. reduced neuronal excitability, increased AP amplitude, diminished inward and outward ionic current densities, while only decreasing inward conductance. Numerous ion channels regulate AP electrogenesis and so it is possible that the same ion channels responsible for the higher rheobase observed in Prdm12-AAV neurons also account for the increased AP amplitude, but this remains to be tested as no pharmacological testing was conducted to identify how the activity of individual ion channels contributed to changes observed. For example, the reduced voltage-gated outward current peak amplitude observed could slow the initiation of the repolarization phase i.e. a decrease in early-activating outward K^+^ currents, which could in turn account for increase in AP amplitude^33^. The substantial attenuation of voltage-gated inward currents and associated conductance, predominantly mediated by voltage-gated Na^+^ channels, could impair the neuron’s capacity to achieve the threshold potential required for action potential initiation^34^. Concurrently, voltage-gated outward currents, albeit diminished, could maintain a hyperpolarizing influence on the membrane potential. This ionic imbalance could effectively elevate the AP threshold, requiring stronger depolarizing stimuli to elicit neuronal firing. Further studies are required to identify changes caused by Prdm12 overexpression and thus might contribute to the phenomena observed.

Several studies have investigated the impact of Prdm12 loss on neuronal excitability, yielding contrasting results^6,7^. In a developmental Prdm12-ablation model, neurons exhibited reduced excitability, evidenced by a significant increase in rheobase, the minimal current required to evoke an action potential^6^. However, this effect was not observed when *Prdm12* was ablated in adult DRG neurons, where rheobase remained unchanged between control and Prdm12 iCKO neurons. In contrast, a later study found that Prdm12 iCKO in adult neurons decreased rheobase, indicating increased excitability in the absence of Prdm12^7^. Numerous reasons could account for differences in the neuronal excitability reported by different groups, including the different strains, knockout strategy, and recording conditions/parameters. Both Latragna and colleagues and Kokotovic and colleagues used tamoxifen-inducible *AdvillinCreERT2; Prdm12fl/fl* mice targeting somatosensory neurons. Both studies used mice on a C57BL/6 background, however, experiments were conducted using a non-defined mix of males and females.

Our observation of a CIP-associated gene potentially exhibiting a comparable phenotype in GOF (this work in adult sensory neurons) and LOF^6^ (Kokotovic and colleagues in developmental sensory neurons) and leading to a decrease in neuronal excitability is not the first to be described. A previous study has presented a GOF mutation p. Leu811Pro alteration in *SCN11A*, that encodes the voltage-gated Na^+^ channel Na_V_1.9, which is highly expressed in nociceptive neurons. Both *SCN11A* LOF *and* GOF mutations led to a decrease in neuronal excitability, and hence CIP^35,36^. Another study has shown that this GOF mutation also leads to significant reduction of the capsaicin-evoked release of calcitonin gene-related peptide indicating reduced neurogenic inflammation^37^. The finding of a GOF mechanism related to this ion channel (*SCN11A*) contrasts with previously described pain-related voltage-gated Na^+^ channel disorders. For example, GOF mutations in *SCN9A*, which encodes Na_V_1*.7*, lead to painful conditions like familial primary erythromelalgia, which contrast to an inability to experience pain in LOF/hypofunctional alterations in the same gene^38–42^. Leipold et al., explained that the impaired pain perception in *SCN11A* GOF mice and humans could be caused by a process in which increased activity of Na_V_1.9 around the resting membrane potential leads to depolarizing block.

Further investigation is required to delineate the mechanisms by which Prdm12 influences neuronal excitability; changes in voltage-gated ionic conductances, leading to altered rheobase, appear to be key. These findings position Prdm12 as a promising target for analgesic intervention, particularly given that its overexpression attenuates inflammatory pain. Future studies should aim to resolve the molecular pathways through which Prdm12 differentially regulates nociceptive neuron sensitivity across distinct pain modalities and other preclinical models of pain.

## 7 Acknowledgments

## Author contributions

Conceptualization: M. Dannawi, E.J. Bellefroid and E. St. J. Smith investigation: M. Dannawi, L. A. Pattison; data analysis: M. Dannawi, L. A. Pattison, A. Cloake; writing: M. Dannawi and E. St. J. Smith; supervision: E.J. Bellefroid and E. St. J. Smith; funding acquisition: E.J. Bellefroid and E. St. J. Smith.

## 8 Figure legends

**Figure S1:**
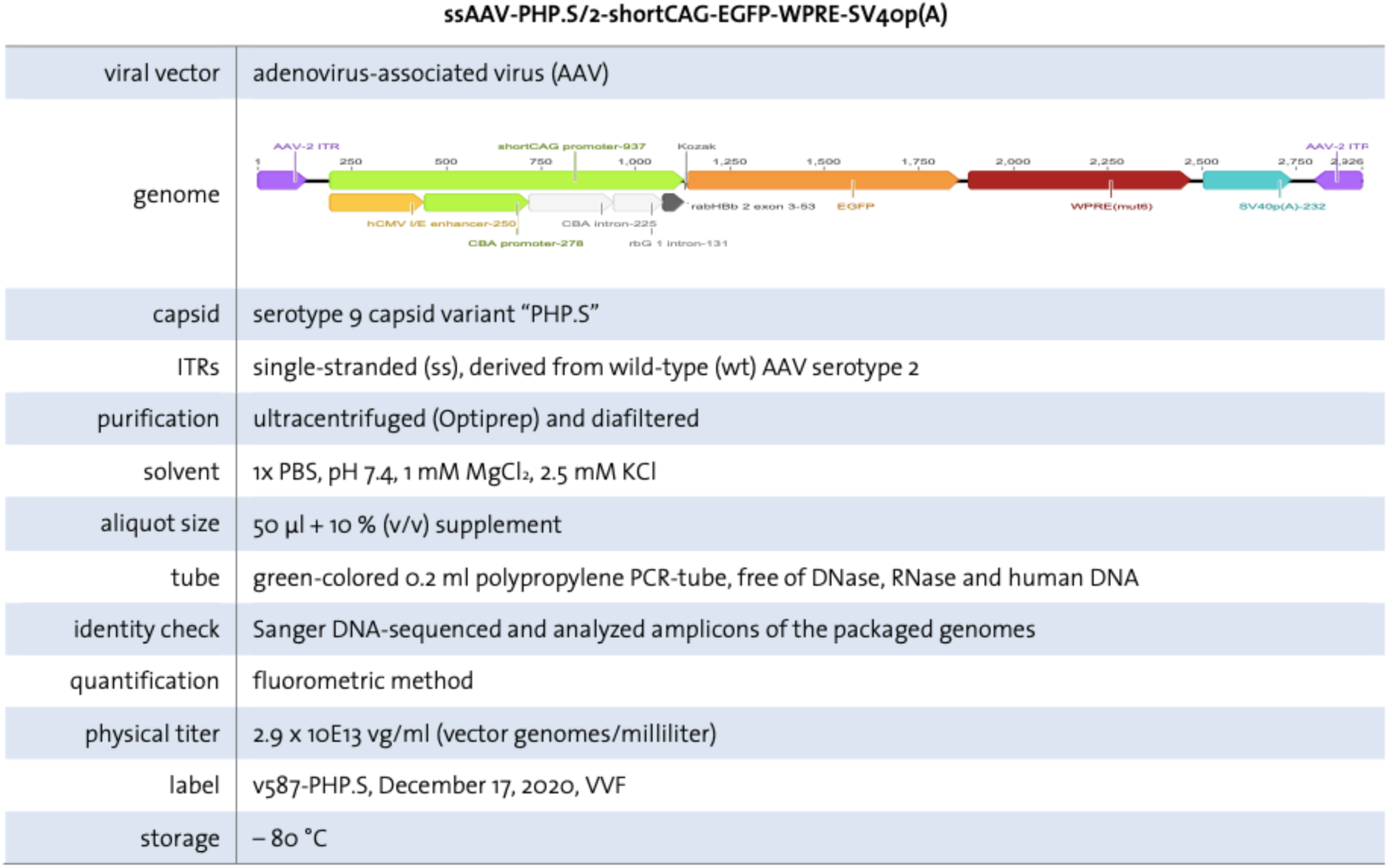
Control-AAV.

**Figure S2:**
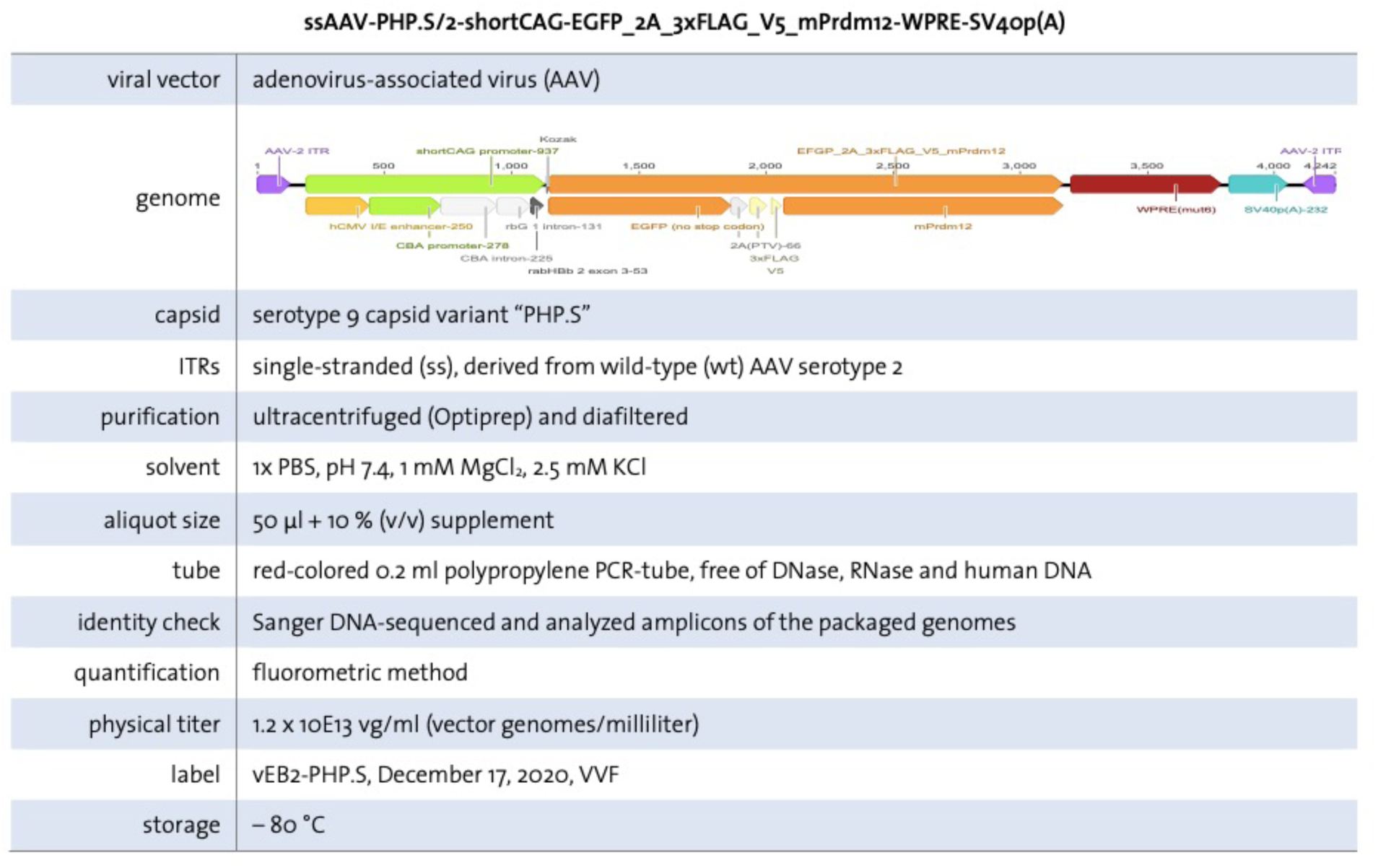
Prdm12-AAV.

**Figure S3:**
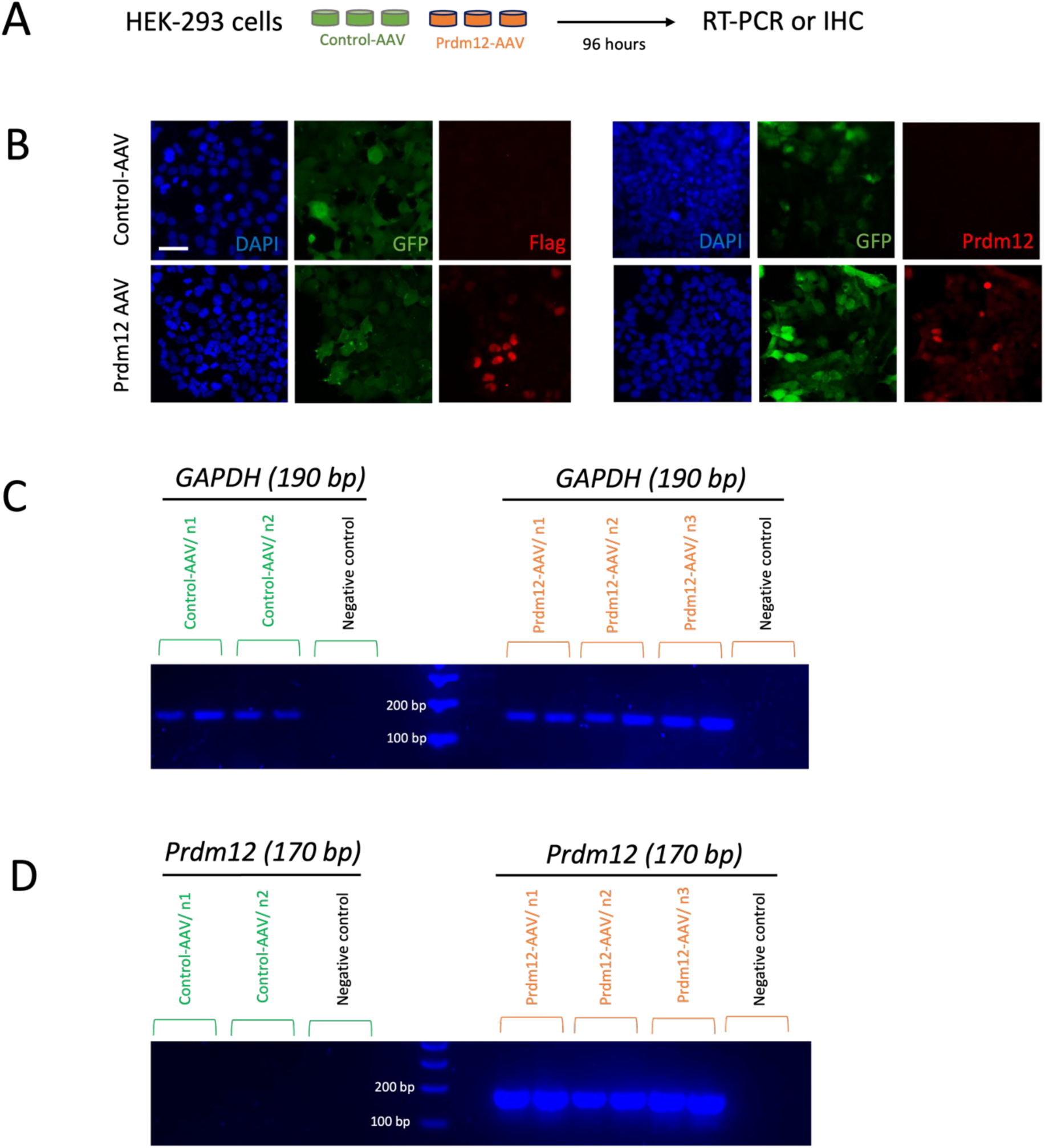
AAV transduction efficiency in HEK-293 cells that normally do not express Prdm12. (A) Scheme of the experimental design (B) immunostaining of HEK cells following 96 hours of AAV incubation comparing GFP (green), Prdm12 (red) and Flag (red) between Control-AAV and Prdm12-AAV conditions. scale bar: 100 μm (C,D) RT-PCR of *GAPDH* (C) and *Prdm12* (D) gene expression comparing *Prdm12* expression between Control-AAV and Prdm12-AAV transduced HEK-293 cells, showing *Prdm12* expression in Prdm12-AAV cells but no expression in Control-AAV cells.

**Figure S4:**
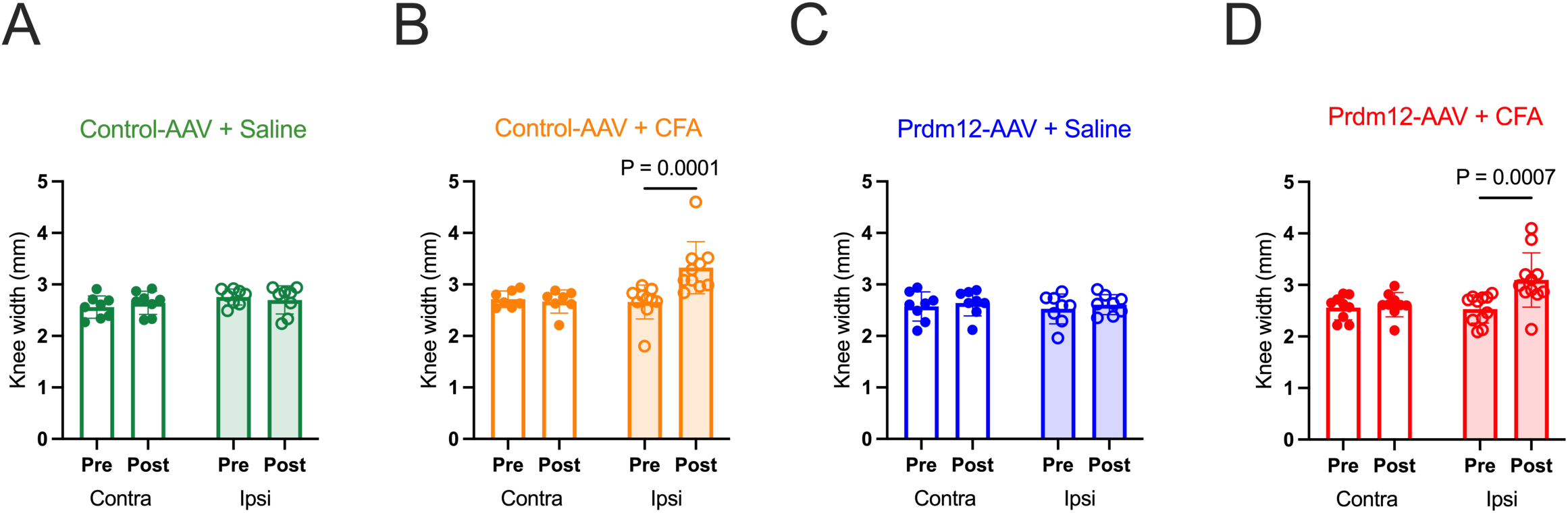
CFA-induced Inflammation reflected by increase in knee width of ipsilateral side. Knee width measurements in Control-AAV and Prdm12-AAV injected mice, comparing pre– and 24 hours post-injection

**Figure S5.**
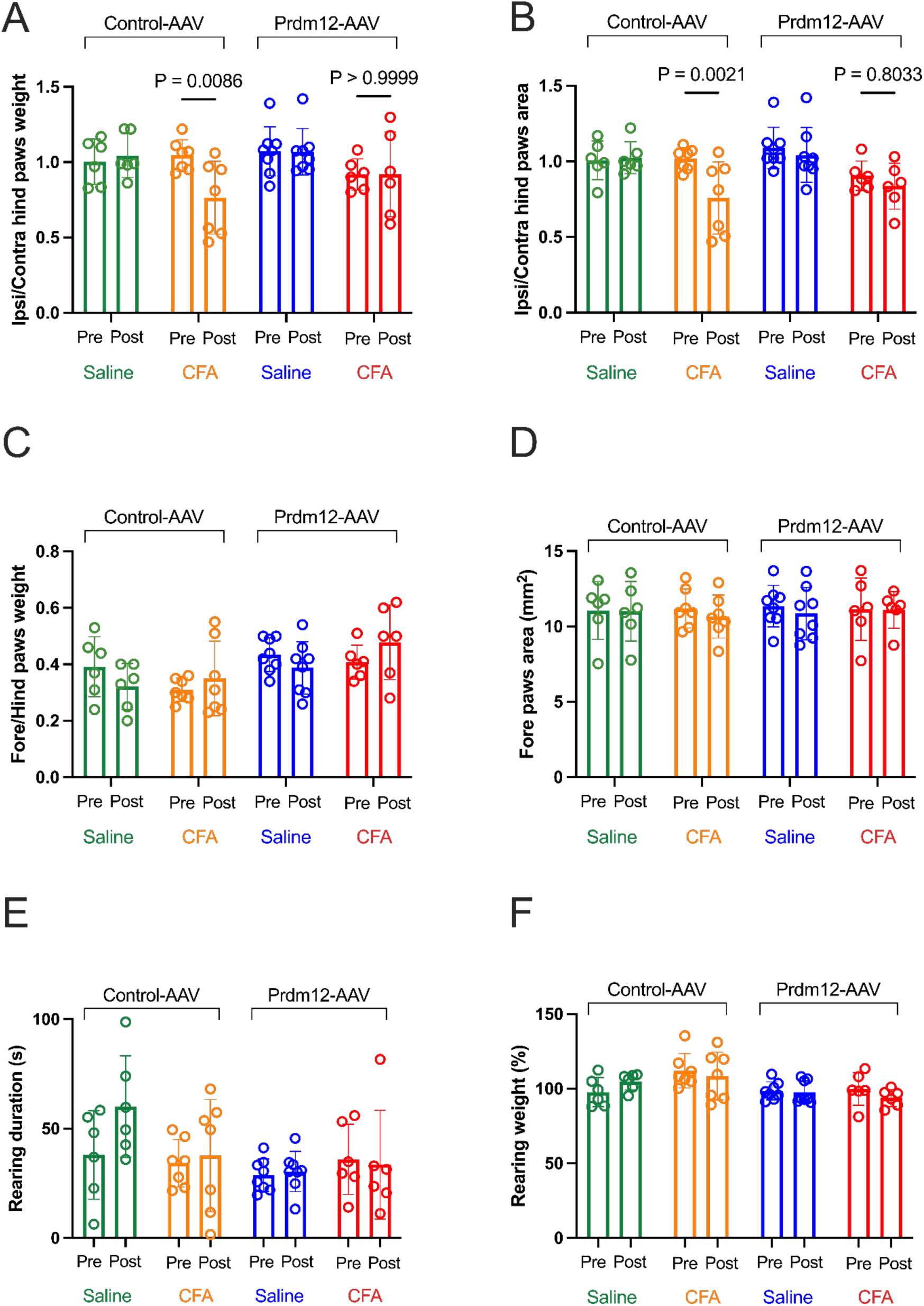
(related to figure 5): Dynamic weight bearing recordings of paws comparing the CFA-injected ipsilateral side vs Saline-injected contralateral side pre and 24 hours post CFA/Saline injection. (A) Weight of ipsilateral hind paw/Contralateral hind paw (B) Ipsi/ Contra hind paws occupied area (C) Weight of fore paw/ hind paw (D) Fore paws occupied area (E) Rearing durations, time spent standing only on hind paws in an upright position (F) Rearing weight when standing on both hind paws. Control-AAV+Saline n=6; Control-AAV+CFA n=7; Prdm12-AAV+Saline n=8; Prdm12-AAV+CFA n=6 recorded over 2 independent experiments (AAV injections) Data represented are mean ± SEM Two-way anova test, followed by Bonferroni test was used

